# sMode of action of the antimicrobial peptide Mel4 is independent of *Staphylococcus aureus* cell membrane permeability

**DOI:** 10.1101/603712

**Authors:** Muhammad Yasir, Debarun Dutta, Mark D.P. Willcox

## Abstract

Mel4 is a novel cationic peptide with potent activity against Gram-positive bacteria. The current study examined the anti-staphylococcal mechanism of action of Mel4 and its precursor peptide melimine. The interaction of peptides with lipoteichoic acid (LTA) and with the cytoplasmic membrane using DiSC(3)-5, Sytox green, Syto-9 and PI dyes were studied. Release of ATP and DNA/RNA from cells exposed to the peptides were determined. Bacteriolysis and autolysin-activated cell death were determined by measuring decreases in OD_620nm_ and killing of *Micrococcus luteus* cells by cell-free media. Both peptides bound to LTA and rapidly dissipated the membrane potential (within 30 seconds) without affecting bacterial viability. Disturbance of the membrane potential was followed by the release of ATP (50% of total cellular ATP) by melimine and by Mel4 (20%) after 2 minutes exposure (*p*<0.001). Mel4 resulted in staphylococcal cells taking up PI with 3.9% cells predominantly stained after 150 min exposure, whereas melimine showed 34% staining. Unlike melimine, Mel4 did not release DNA/RNA. Cell-free media from Mel4 treated cells hydrolysed peptidoglycan and produced greater zones of inhibition against *M. luteus* lawn than melimine treated samples. These findings suggest that pore formation is unlikely to be involved in Mel4-mediated membrane destabilization for Staphylococcci, since there was no significant Mel4-induced PI staining and DNA/RNA leakage. It is likely that the *S. aureus* killing mechanism of Mel4 involves the release of autolysins followed by cell death. Whereas, membrane interaction is the primary bactericidal activity of melimine, which includes membrane depolarisation, pore formation, release of cellular contents leading to cell death.

This work is original, has not been published and is not being considered for publication elsewhere. Part of this manuscript has been presented as a poster presentation in Gordon Research Conference Italy in 2019. There are no conflicts of interest for any of the authors that could have influenced the results of this work. Prof. Mark Willcox holds the patent the for the melimine peptide.

## INTRODUCTION

*S. aureus* is a major cause of infections in both health care and community settings which can produce high levels of mortality and have a high economic burden on society [1, 2]. *S. aureus* can reside on the skin and in the nasal cavity of 20–50% humans [3, 4] and this poses a risk for subsequent infections [5]. *S. aureus* is also a major cause of infections of medical devices and can cause 30–40% bacteraemia, surgical wound and implant-related infections [4, 6]. Methicillin-resistant *S. aureus* (MRSA) now causes over 50% of skin and soft tissue infections [7]. The mortality rate of *S. aureus* bacteraemia can reach upto 40% [8]. *S. aureus* associated infections are difficult to treat with currently available antibiotics [9] partly due to the increase in MRSA which are often very resistant to many different classes of antibiotics [6, 10]. To overcome these problems, new antimicrobials are needed which have unique modes of action and limited potential for resistance development.

Antimicrobial peptides (AMPs) exhibit broad spectrum antimicrobial activity against a wide range of microorganisms including bacteria, fungi, parasites and enveloped viruses at low concentrations [11-13]. AMPs are usually cationic in nature and have a varying number (from five to over a hundred) of amino acids. AMPs possess multiple modes of action, rapid bacterial killing kinetics and little toxicity toward human cells [14, 15]. Owing to the fact that these molecules exhibit multifaceted modes of action and rapidly kill bacteria, development of resistance in bacteria against AMPs is relatively rare [16, 17]. The mechanism of action of AMPs is believed to start by interacting with the negatively charged lipoteichoic acid (LTA) or teichoic acid of Gram-positive bacteria [18], through the negatively charge phosphate groups in LTA [19] [20].

The interaction with LTA is then believed to facilitate penetration of AMPs through the thick peptidoglycan layer, perhaps by the LTA acting as ladder, allowing the AMPs to reach and act on the cytoplasmic membrane [18]. AMPs disrupt phospholipid lipid bilayers of bacterial membranes by forming pores by various mechanisms called “barrel stave” or “toroidal pore” or through disintegration of lipids via the “carpet model” [21]. Cytoplasmic membrane collapse can result in leakage of cellular contents such as potassium ions, ATP and DNA/RNA which in turn may lead to cell death [22, 23]. Some AMPs translocate across the cell membrane and inhibit DNA/RNA or protein synthesis [24, 25]. AMPs can also kill Gram positive bacteria by activating cell wall bound autolytic enzymes known as autolysins [26]. LTAs bind to autolysins in bacterial cell walls and control their activity. AMPs can disturb the regulatory function ofLTA resulting in unregulated autolysin activity which then acts on the peptidoglycan chains and peptide bridges of murein [27].

Melimine is a cationic hybrid peptide of melittin and protamine [28]. It shows broad spectrum antimicrobial activity against Gram-negative and Gram-positive bacteria (including MRSA), fungi and protozoan such as *Acanthamoeba* without inducing resistance in bacteria [28, 29]. Melimine destroys the membrane potential of *S. aureus* [30]. Melimine, covalently bound to polymers such as contact lenses, can inhibit initial bacterial attachment and kill microbes [28, 29]. A shorter sequence of melimine, called Mel4, has a relatively low minimum inhibitory concentration against *S. aureus* (52.3 μM) and kills *S. aureus* when immobilized on surfaces [31]. It is non-cytotoxic to mammalian cells *in vitro* [28, 29]. Mel4 has improved ocular compatibility compared to melimine when bound to contact lenses [32]. Mel4 lacks the amino acids tryptophan, leucine and isoleucine that are present in the sequence of melimine. Tryptophan is a highly lipophilic amino acid [33] and its presence is often an important part of the activity to AMPs [34-36]. Leucine and isoleucine are hydrophobic residues which promote strong α-helix formation in AMPs which in turn results a higher degree of membrane disruption [37, 38]. Although various studies have shown that Mel4 has high bactericidal activity aginst staphylococci, the mode of action of Mel4 is not yet understood. Given the shorter peptide length and the unique amino acid sequence compared to melimine, an investigation of the killing mechanism of Mel4 against Gram-positive bacteria is neccessary. Hence, this study was designed to evaluate the mode of action of Mel4 in comparison with melimine against *S. aureus.*

## MATERIALS AND METHODS

### Synthesis of peptides

Melimine (amino acid sequence: TLISWIKNKRKQRPRVSRRRRRRGGRRRR; molecular mass 3786.6) and Mel4 (KNKRKRRRRRRGGRRRR; 2347.8) were synthesized by conventional solid-phase peptide protocols [39, 40] and procured from the Auspep Peptide Company (Tullamarine, Victoria, Australia). The purity of the purchased peptides was ≥90%.

### Bacterial strains and growth conditions

*S. aureus* 31 (contact lens induced peripheral ulcer isolate) [41], *S. aureus* 38 (microbial keratitis isolate) [42], and the type strain of *S. aureus,* ATCC 6538 (isolated from a “human lesion” according to the ATCC website) [43] were used in the current study. Bacteria were grown from frozen stocks overnight at 37°C to mid-log phase in Tryptic Soy Broth (TSB; Oxoid, Basingstoke, UK) and cells were harvested following washing with phosphate buffer saline (PBS, NaCl 8 g/L, KCl 0.2 g/L, Na_2_ HPO_4_ 1.4 g/L, KH_2_ PO_4_ 0.24 g/L; pH 7.4) then diluted in the PBS containing 1/1000 TSB to OD_600nm_ 0.05-0.06 (1× 10^7^ colony forming units (CFU)/ml confirmed upon retrospective plate counts on TS agar (Oxoid)). Cells prepared this way were used in most experiments except for assessing the minimum inhibitory and bactericidal concentrations, measuring the release of DiSC3-5 from cells, and the autolytic activity assay.

### Minimum inhibitory and minimum bactericidal concentration

The minimum inhibitory and minimum bactericidal concentrations of melimine and Mel4 were determined for all strains using a modified version of the Clinical Laboratory and Standard Institute (CLSI) broth microdilution method as described previously [44]. Bacterial suspensions were prepared to a final concentration of 5×10^5^ CFU ml^-1^ in Muller Hinton Broth (MHB, Oxoid) and added to wells in a sterile 96-well microtiter plate containing two-fold dilutions of each peptide with 0.01% v/v acetic acid (Sigma Aldrich, St Louis, MO, USA) and 0.2% w/v bovine serum albumin (Sigma Aldrich). The plate was incubated at 37°C for 18 h-24 h. The MIC was set as the lowest concentration of peptides that reduced bacterial growth by ≥ 90% while the MBC was set as the lowest concentration of peptides that reduced bacterial growth by >99.99% after enumeration of viable bacteria by plate counts compared to bacteria grown without either AMP.

### Interaction with LTA

Two experiments, bacterial growth in the presence of LTA and the amount of LTA in solution measured by ELISA, were performed to determine the interaction of melimine and Mel4 with LTA. Growth of *S. aureus* 38 was conducted in the presence of melimine or Mel4 following a method of Yang and Yousef [45] with slight modifications. *S. aureus* LTA (Sigma Aldrich; 100 µg/ml) was dissolved in MHB in wells of a 96-well microtiter plate containing *S. aureus* (5 ×10^5^ CFU/ml) followed by addition of 100 µl of melimine or Mel4 to achieve their MICs. The OD_600nm_ was determined over time during incubation at 37° C. A growth curve was constructed over 12 h. Controls were bacteria grown in the presence of LTA alone and bacteria grown in MHB alone (a measure of maximum bacterial growth).

A competitive ELISA using kit (My BioSource, San Diego, CA, USA) was performed to evaluate whether melimine and Mel4 could bind and neutralize LTA of *S. aureus*. LTA (200 ng/ml) was dissolved in endotoxin free water (Sigma Aldrich) with melimine or Mel4 at their MICs or MBCs as final concentrations and incubated for 1-2 h at 37 °C. Interaction of LTA with melimine and Mel4 was assessed as an increase in OD_450nm_ compared to negative control without peptides according to manufacturer instructions.

### Cytoplasmic membrane disruption

The effect of melimine and Mel4 on the cytoplasmic membrane of *S. aureus* was assessed in three experiments. To examine membrane depolarisation, the release of the membrane potential sensitive dye DiSC3-5 was measured. Sytox Green and Propidium Iodide (PI) dyes were used to determine whether the peptides could damage the cytoplasmic membranes and allow the stains to penetrate and interact to intracellular nucleic acids. Sytox Green has a molecular mass of 213.954 g/mol and a topological polar surface area of 28.7 A^2^ [46] whereas PI has a molecular mass of 668.087 g/mol and a topological polar surface area of 55.9 A^2^ [47]. Penetration of these two dyes through compromised membrane occurs through pores and differences may indicate different pore structures/sizes created by peptides.

Cytoplasmic membrane depolarization was performed as described by Rasul *et al.,* [30] with slight changes. Aliquots 100 µl of DiSC3-5 (Sigma Aldrich, St Louis, MO, USA) labelled bacteria in HEPES [30] and 100 µl of melimine and Mel4 at their MICs and MBCs (as final concentrations) were added to wells of 96 well microtiter plates. Increases in fluorescence were determined at 30 second intervals at an excitation wavelength of 622 nm and an emission wavelength of 670 nm over five minutes. Aliquots of bacteria were withdrawn over the time course and serially diluted in Dey-Engley (D/E neutralizing broth (Remel, Lenexa, KS, USA)) then plated on Tryptic Soy Agar (Oxoid, Basingstoke, UK) containing phosphatidylcholine (0.7 g /L) and Tween 80 (5ml/L) then incubated at 37 °C overnight. The number of live bacteria were enumerated and expressed as CFU/ml. A positive control of dimethyl sulfoxide (DMSO) (Merck, Billerica, MA, USA) (20% v/v) in HEPES (100 µl) was used to disrupt the cytoplasmic membrane potential of bacteria [48].

Sytox green assay was performed following the established protocol of Li *et al.,*[49] with slight modifications. Briefly, bacterial cells in PBS (100 µl) were dispensed into wells of 96-well plates along with 5µM Sytox green (Invitrogen, Eugene, Oregon, USA). Plates were incubated for 15 minutes in the dark at room temperature and then 100 µl of melimine or Mel4 were added so as to achieve their MICs and MBCs. Fluorescence was measured spectrophotometrically (at an excitation wavelength of 480 nm and an emission wavelength of 522 nm) every 5 minutes up to 30 minutes, and then after 150 minutes. Triton X-100 1% (v/v) (Sigma Aldrich, St Louis, MO, USA) in PBS with 1/1000 TSB (100 µl) was used as positive control to disrupt the cytoplasmic membrane of bacteria.

Flow cytometry was used to determine the ability of melimine and Mel4 to permeabilize the cytoplasmic membrane of *S. aureus* 38 only. Bacteria were incubated simultaneously with SYTO9 (7.5 µM) and PI (30 µM; Invitrogen, Eugene, Oregon, USA) and incubated at room temperature for 15 min. Fluorescence intensities were recorded with a LSRFortessa SORP Flow cytometer after addition of melimine or Mel4 at their MIC at different time points. The wavelength of green fluorescence was (525/550 nm) for SYTO9 and a red fluorescence (610/20 nm) for PI [50]. Data were acquired and analyzed using Flowjo software (version 10.5.0, Oregon, USA). A minimum 20000 events were recorded for each sample.

### Leakage of intracellular contents

Two assays, ATP release and loss of DNA/RNA. were performed to determine the leakage of intracellular contents. Bacterial suspensions (100 µl) were added to melimine and Mel4 at their MICs and MBCs and incubated at 37°C for 10 minutes. Samples were withdrawn at two minutes intervals and centrifuged at 9000 × g for five minutes, and the supernatant removed and kept on ice until further analysis. For determination of total ATP, the bacterial pellet was resuspended in boiling 100 mM Tris, 4 mM EDTA pH (7.4) and further incubated for 2 mins at 100 °C to lyse all the cells. The lysed cells were centrifuged at 9600 × g for two minutes and the supernatant was kept on ice until further use [51]. Subsequently, both total and released ATP were determined using an ATP bioluminescence kit (Invitrogen, Eugene, Oregon, USA) according to the manufacturer’s instructions.

The loss of DNA/RNA was determined by following a method of Carson *et al.,* [52] with slight modifications. Bacteria (100 µl) were incubated at 37 °C with melimine and Mel4 at their MICs and MBCs. Samples were withdrawn at 5 minutes intervals, diluted (1:10) and filtered through 0.22 µm pores (Merck, Tullagreen, Ireland). The OD_260nm_ of the filtrates was measured in a UV-star plate (Greiner Bio-one GmbH, Frickenhausen, Germany). The results were expressed as the ratio to the initial OD_260nm_.

### Lysis of bacteria

The bacteria-lytic potential of the two peptides was evaluated using two different bacterial inoculums 1× 10^8^ CFU/ml (OD_660_ 0.1) and 3 × 10^10^ CFU/ml. The smaller inoculum size was used to see whether OD_620nm_ was measurable at the concentration of cells used in all other assays. However, no measurable optical density change at 660nm was obtained, and so a larger inoculum of 3 × 10^10^ CFU/ml was used. The larger inoculum was obtained by adjusting OD_620nm_ to 0.3, and bacterial numbers (CFU/ml) were confirmed upon retrospective plate count. Melimine and Mel4 were added at their MICs and MBCs. Bacterial cultures were immediately mixed and then diluted 1:1000 in TSB. The OD_620nm_ was measured and additional readings were taken at 30, 60, 90, 120 minutes, 6.5 and 24 h after incubating at room temperature. Peptides at their respective concentrations in 1/1000 TSB were used as blanks. The results were recorded as a ratio of OD_620nm_ at each time point compared to the OD _620nm_ at 0 minutes (in percentage) [52].

### Time kill assay

Aliquots of bacterial cells (100 µl; 1× 10^7^ CFU/ml) were added to melimine or Mel4 at their MICs and MBCs (as final concentration) in 96 well plates and incubated at 37°C with gentle shaking. Aliquots (100 µl) were withdrawn at specified time points, serially diluted into D/E neutralizing broth and plated onto Tryptic Soy agar containing phosphatidylcholine (0.7 g /L) and Tween 80 (5ml/L). Bacterial colonies were counted after 24 h of incubation at 37 °C. Viable bacteria were reported as log_10_ CFU/ml. A control of bacterial suspensions without peptide was performed under the same conditions. The assay was performed in triplicate.

### Release of autolysins

The presence of autolysins in supernatants of strains (*S. aureus* 31 and *S. aureus* 38) grown in the presence of melimine and Mel4 was determined spectrophotometrically by the ability of the supernatants to decrease the optical density of *Micrococcus luteus* peptidoglycan (PGN; Sigma Aldrich, St Louis, MO, USA) following a published method [53] with minor modifications. Bacteria were grown in MHB and harvested by washing three times with PBS. Bacteria were resuspended in the same PBS to an OD_660nm_ of 0.1 (1× 10^8^ CFU/ml). The bacterial suspension (100 µl) was incubated at 4X MIC of melimine or Mel4 for 5 h at 37 °C. Following centrifugation at 2000 × *g* for 10 minutes at 4 °C, the supernatant was recovered and filtered through 0.2 µm filter (Merck, Tullagreen, Ireland). The filtrate was saturated with 75% ammonium sulphate at 4 °C with continuous mixing for 1 h. This mixture was centrifuged at 12000 × *g* at 4 °C for 30 min, the supernatant was discarded, and the pellet was resuspended in 0.1 M sodium acetate buffer (pH. 6) and dialysed against four changes of buffer using Amicon ultra centrifugal filters (Merck, Millipore, Tullagreen, Carrigtwohill, IRL). The retentate was recovered and incubated with 100 µl of *Micrococcus luteus* peptidoglycan (1 mg/ml) at room temperature for 3 h. OD_570 nm_ was initially determined at 0 min and then every 30 min intervals until 3 h. Results were presented as a ratio of OD_570 nm_ at each time point versus OD_570nm_ at 0 minutes (in percentage). The retentate from the above assay was also used to detect the autolytic activity against a lawn of *Micrococcus lysodeikticus* ATCC 4698. Briefly, 10 μL of each retentate from the above assay was spotted onto a pre-seeded lawn of *M. lysodeikticus* (10^8^ CFU/ml) on nutrient agar. Following incubation for 24 h at 37 °C, the size of the zone of inhibition was measured. Lysozyme (5 mg/ml) used as a positive control for maximum growth inhibition. Melimine or Mel4 dissolved in PBS at 4X MIC was treated in exactly the same way but in the absence of bacteria, to test whether any of the inhibitory activity in both assays (PGN and lawn of bacteria) was due to the presence of these peptides.

### Statistical analyses

Statistical analyses were performed using GraphPad Prism 7.02 software (GraphPad Software, La Jolla, CA, USA). The effect of the different concentrations of peptides in each assay compared to controls was analysed using two-way ANOVA with Tukey’s test of multiple comparisons. Correlations between release of extracellular ATP and bacterial death were examined using Pearson correlation test. Statistical significance was set as *p*<0.05.

## RESULTS

### Inhibitory concentrations of the two peptides

The MICs and MBCs for melimine or Mel4 against *S. aureus* are shown in Table 1. Both melimine and Mel4 were most active against *S. aureus* ATCC 6538. Melimine had lower MIC and MBC compared to Mel4 for all strains.

**Table 1.**
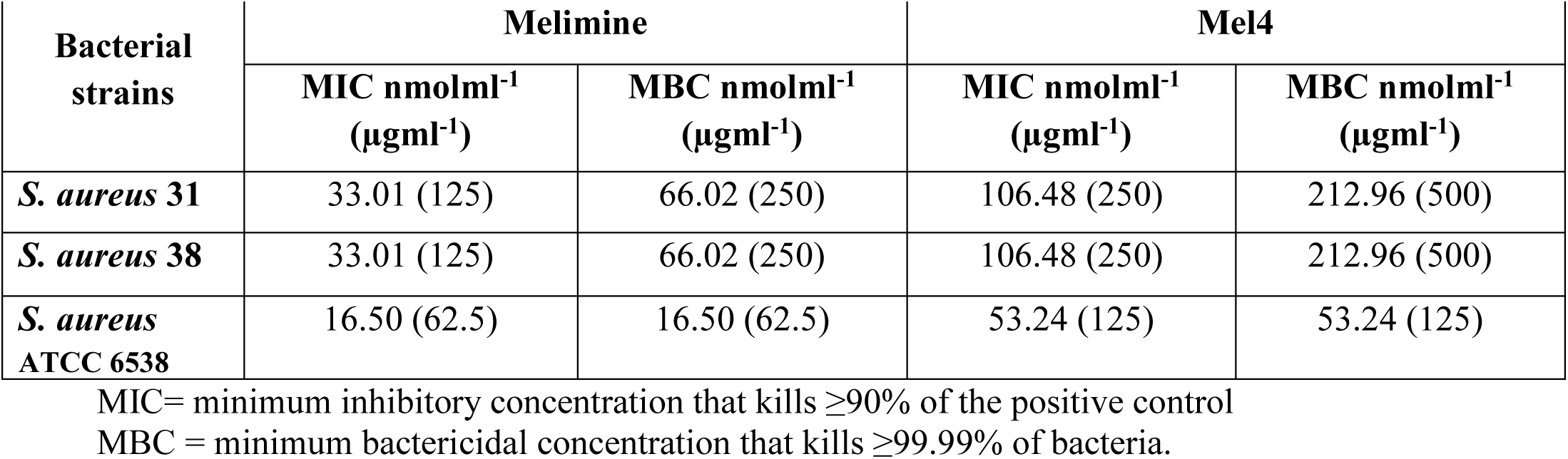
MIC and MBC values of melimine and Mel4 against *S. aureus*

| Bacterial strains | Melimine |  | Mel4 |  |
| --- | --- | --- | --- | --- |
|  | MIC nmolml<sup>-1</sup><br>(µgml<sup>-1</sup>) | MBC nmolml<sup>-1</sup><br>(µgml<sup>-1</sup>) | MIC nmolml<sup>-1</sup><br>(µgml<sup>-1</sup>) | MBC nmolml<sup>-1</sup><br>(µgml<sup>-1</sup>) |
| *S. aureus* 31 | 33.01 (125) | 66.02 (250) | 106.48 (250) | 212.96 (500) |
| *S. aureus* 38 | 33.01 (125) | 66.02 (250) | 106.48 (250) | 212.96 (500) |
| *S. aureus* ATCC 6538 | 16.50 (62.5) | 16.50 (62.5) | 53.24 (125) | 53.24 (125) |

MIC= minimum inhibitory concentration that kills ≥90% of the positive control
MBC = minimum bactericidal concentration that kills ≥99.99% of bacteria.

In all subsequent experiments data of the assays for *S. aureu*s 38 is presented. Data for all other strains are presented in the Supplementary information (except flow cytometry where only *S. aureus* 38 was used, and autolysins where only *S. aureus* 38 and 31 were used). Both the AMPs displayed similar modes of action against all the strains of *S. aureus.*

### Interaction with LTA

Melimine at its MIC (33.01 nmol/ml) and Mel4 at its MIC (106.48 nmol/ml) inhibited the growth of *S. aureus* (Fig.1) with OD_600 nm_ <0.2 after 14 hours of growth. However, in the presence of LTA *S. aureus* could grow with significant increases in OD_600nm_ starting after 6 h (*p*<0.001; Fig. 1) with both peptides. The OD_600 nm_ of bacteria with LTA-melimine or LTA-Mel4 was not different at 6 h (*p*=0.601). The presence of LTA alone did not affect the growth of *S. aureus*. In the ELISA assay at their MICs, melimine neutralized 1.2±0.1 ng LTA/nmol and Mel4 neutralized 1.1±0.1 ng LTA/nmol (no significant difference). Similarly, at their bactericidal concentrations they neutralized 0.8±0.1 ng/nmol and 0.6±0.1 ng/nmol respectively (no significant difference).

### Membrane disruption

Melimine depolarized the cytoplasmic membrane of *S. aureus* in a predominantly concentration independent manner. An increase in fluorescence intensity was detected as early 30 seconds after addition of either peptide to all strains of *S. aureus.* The increase in fluorescence was continuous until at 180 sec and became constant after this time until the last observation at 300 sec (Fig. 2). The fluorescein intensity was not statically different at the MIC and MBC of melimine (*p*>0.999). However, at its MBC Mel4 yielded a significantly higher fluorescence than at its MIC at 60 to 90 sec (*p*<0.05) and after this time the degree of depolarization was similar for both the concentrations. This depolarization of the cytoplasmic membrane was not directly associated with the bacterial death as only < 0.5 log_10_ bacteria were affected with both the peptides at either concentration (Fig. 2).

The fluorescence of Sytox green increased over time only with melimine and no fluorescence was detected with Mel4. Melimine at its MIC or MBC permeabilized the cell membrane and the fluorescence increased significantly after 10 minutes of exposure compared to buffer treated controls (*p*<0.026) (Fig. 3). The intensity of fluorescence gradually increased over 150 minutes for both the concentrations of melimine against all strains, but no significant difference was observed between the MIC and MBC (*p*>0.999). Treatment with the positive control Triton-X 100 resulted emission of higher Sytox green fluorescence compared to either peptides within 10 minutes (*p*<0.001).

The membrane damaging effect of these peptides was also assessed with *S. aureus* 38 only by flow cytometry in the presence of the DNA intercalating dye PI and the membrane permeable dye Syto-9 (Fig. 4). After 30 minutes exposure to melimine at its MIC, 22.2% of cells had high levels of Syto-9 staining but low levels of PI, 66.31% of cells had high levels of both Syto-9 and PI staining and 10.1% of cells had high levels of PI but low levels of Syto-9 staining. The total percentage of cells with high levels of PI staining after 30 minutes exposure to melimine was 76.41%. Whilst after 150 minutes the total number of PI stained cells remained approximately the same at 80.4%, the proportions had changed with 34.2% of cells having high levels of PI staining and low levels of Syto-9 staining and 46.2% having high levels of both Syto-9 and PI staining. The time course of staining with Mel4 at its MIC was different to melimine. After 30 minutes incubation, 55.3 % of cells had high levels of Syto-9 staining but low levels of PI, 32.8% of cells had high levels of both Syto-9 and PI staining and 3.58% of cells had high levels of PI but low levels of Syto-9 staining. The total percentage of cells with high levels of PI staining after 30 minutes exposure to melimine was 36.4%. After 150 minutes the total number of PI stained cells remained approximately the same at 41.6%, as did the relative proportions with 3.91% of cells having high levels of PI staining and low levels of Syto-9 staining and 37.7% having high levels of both Syto-9 and PI staining. However, when looking at the flow cytometry readout (Fig. 4) there did appear to be a general increase in the percentage of double stained cells, but the gating did not bin cells with sufficiently fine detail. There was a difference in the kinetics of cell death with the positive control Triton-X 100 (1%), which after 30 minutes stained 16.34% cells with high PI and increased up to 24.94 % after 150 min incubation. With buffer treated (negative control) cells > 86% were stained with high levels of Syto-9 and low levels of PI even after 150 minutes incubation (Fig. 4).

### The release of cytoplasmic contents

Melimine at its MIC or MBC released 49% and 51% of total cellular ATP respectively after 2 minutes of exposure (*p*<0.001) (Fig. 5) and this amount of ATP did not increase after the initial burst at 2 minutes. Within the first two minutes, melimine decreased live bacteria by approximately 1.4 log_10_ at its MIC and 1.5 at its MBC, although this was not statistically different to each other and there was no further reduction in bacterial numbers to the end of the assay period (10 minutes). Compared with melimine, Mel4 released significantly (*p*>0.999) less ATP at its MIC (19%) and MBC (21%) after 2 min (Fig. 5), with no increase up to 10 minutes. There were < 0.5 log_10_ viable bacteria after 2 minutes incubation with Mel4 at its MIC or MBC, with no further cell death over the remaining 8 minutes of the assay. Incubation with melimine released higher amounts of ATP and produced more bacterial death (*p*<0.001) than Mel4 at the MICs or MBCs.

Release of DNA/RNA (260nm absorbing materials) is shown in Fig. 6. Incubation with melimine resulted in release of DNA/RNA from *S. aureus* in time dependent but dose independent manner. Although increase in optical density first happened at 5 minutes with melimine it was not significant (*p*≥0.083). A significant increase in the optical density occurred after 10 minutes exposure with increases of 1.7 times and 2.0 times at MIC and MBC respectively (*p*<0.001). Moreover, optical density increased 4.8 times at MIC and 5.1 times at MBC after 150 minutes. On the hand, incubation with Mel4 did not increase optical density at its MICs or MBCs over the course of 150 minutes of the assay (*p*≥0.9945).

### Bacterial lysis

At low bacterial concentrations (1× 10^8^ CFU/ml), both the peptides showed no effect on optical density. This was probably due to bacterial concentration was too low to detect any change in optical density. However, when higher inoculum size of 1× 10^10^ CFU/ml was used and treated with various concentration of peptides, a significant reduction in optical density was observed at 6.5 hours (Fig. 7). At this time point, melimine reduced optical density by 21±1% at its MIC and 31±4% at its MBC compared to buffer treated control (*p*<0.001) (Fig. 7). Melimine lysed more than 42% cells after 24 h. A similar trend was observed for Mel4 which reduced optical density by 22±3 % and 25±2 % at its MICs and MBCs respectively after 6.5 h (Fig. 7). After 24 h of incubation, Mel4 caused lysis of more than 40% cells. Over 24 hours the bacterial-lytic efficiency of both melimine and Mel4 was similar and no significant difference was observed (*p*≥0.851). The optical density of control cells (buffer treated) remained unchanged over the 24 h of the experiment.

The killing kinetics of AMPs was determined by exposing *S. aureus* to their MIC and MBC. Melimine reduced the number of cells by 1.1 log_10_ at its MIC and 1.2 log_10_ CFU/ml at its MBC after 2 h of incubation (Fig. 8). At its MIC, after 2 hours melimine continued to reduce the numbers of live cells, but the rate of reduction diminished after 12 hours. The initial rate of reduction from 0-2 h was 6.6 CFU log_10_ /ml/hour, from 2-12 hours was 5.9 CFU log_10_ /ml/hour and from 12-24 h was 2.9 CFU log_10_ /ml/hour. At its MBC, the initial rate of reduction was 6.7 CFU log_10_ /ml/hour, then from 2-24 h the rate remained constant at 5.7 CFU log_10_ /ml/hour. Mel4 at its MIC did not significantly reduce bacterial viability within the first 2 h. The rate of reduction from 2-12 h was 5.6 CFU log_10_ /ml/hour and 3.8 CFU log_10_ /ml/hour from 12-24 h. At its MBC the rate of reduction was also negligible initially (0-2 h), however, from 2-24 h the rate of reduction increased to 5.5 CFU log_10_ /ml/hour. Killing rate of melimine from 2-24 was significantly higher than Mel4 (*p*<0.001). The bacterial viability remained unaffected in control (buffer treated) over 24 h of the experiment.

### Autolytic activity

Autolytic activity of peptide-treated *S. aureus* supernatants was tested using PGN as the substrate. The supernatant of Mel4 treated *S. aureus* had a higher reduction of OD_570nm_ of PGN suspension than melimine treated or controls (bacteria grown in the absence of peptides) (Fig. 9). The supernatants of Mel4 treated *S. aureus* resulted in a decrease in PGN density of 17±3% which was significantly more compared with melimine treated (8±2%; *p*=0.004) and controls (7±1%, *p*<0.001) at 60 min. The reduction in PGN density caused by melimine or buffer supernatants was not significantly different at each time point (*p*=0.999). The rate of PGN degradation for Mel4 was 0.28% OD_570_ /min up to 60 mins and then there was no more activity from 60-180 minutes. The rate for lysozyme was 0.7% OD_570_ /min within first 30 minutes. Lysozyme produced higher PGN hydrolysis, decreasing OD 21±4% at 30 min. The autolysis of culture supernatants was also examined using a pre-seeded lawn of *Micrococcus lysodeikticus*. Melimine or Mel4 suspended in buffer and treated in the same way as culture supernatants of *S. aureus* did not produce any zone of inhibition, showing that the peptides must either have been destroyed or removed during processing. The supernatant from Mel4 treated cells produced a zone of inhibition of 8±1 mm whereas for melimine and buffer treated cells smaller zones of inhibtion of 5±1 mm were produced. The positive control lysozyme formed largest 12±2 mm zone of inhibition.

## DISCUSSION

The current study demonstrated that the mode of action of *S. aureus* killing by Mel4 and its precursor melimine were different. A previous study had shown that 30 minutes incubation with melimine can distort *S. aureus* cells, induce bleb formation and accummulation of cell debris [30], as well as being able to permeabilise the cytoplasmic membrane of *S. aureus* in a concentration independent manner [30]. The previous study also showed that melimine was only able to produce <1 log_10_ reduction in numbers of *S. aureus* cells when incubated for up to five minutes. The current study demonstrated similar effects, but extended this to demonstrate that melimine could interact with LTA of *S. aureus* and produce pores in the cytoplasmic membrane of *S. aureus* that resulted in leakage of intracellular contents (ATP and DNA/RNA) eventually leading to cell lysis and death. Melimine was also able to liberate a small amount of autolysins from the staphylococcal cells. Whilst Mel4 could also interact with LTA and depolarise the cytoplasmic membrane. It had a weaker effect on producing pores which was evidenced by negligible Sytox green intake by cells, minimum disruption to cytoplasmic membrane compared to melimine and minimum propidium iodide staining. Based on the flow cytometry data it is suggested that it can cause tansient cell membarne permeability. In addition, less ATP was released from Mel4-treated cells and no DNA/RNA was released over 150 minutes. However, both the peptides eventually resulted in cell lysis and death after 24 hours exposure. Mel4-treated cells released greater amount of autolysins compared to melimine-treated ones. It is probable that the main killing mechanism of Mel4 was release of autolysins from LTA in cell walls along with membrane depolarisation, whereas the main killing mechanism for melimine was membrane pore formation together with release of autolysins.

There are substantial differences in the amino acid compositions of Mel4 and melimine which contains 17 and 29 amino acids respectively. The lack of tryptophan in the sequence of Mel4 may be important in the differences in modes of action. Tryptophan (Trp) is known to interact with lipid bilayers, can enhance peptide-membrane interactions and facilitate insertion of peptides into the membrane [34, 54]. Trp and Arginine (Arg) together are known to promote stable and deeper insertion into the cell wall of Gram-positive bacteria [55]. Trp facilitates the insertion of Arg into hydrophobic regions of membranes *via* cation–π interactions causing rapid membrane disruption [34]. May be due to the presence of Trp, melimine can adopt a partial α helix in bio-membrane mimetic environment [30]. The antimicrobial activity of AMPs is often attributed to this helical conformation [56]. Mel4 lacks tryptophan and also has a low hydrophobic moment (0.039), meaning that it is less likely to be attracted within lipid bilayers [57]. Mel4 is unable to interact with lipid spheroids composed of 1-oleoyl-2-hydroxy-sn-glycero-3-phosphocholine (PC 18:1) or tethered lipid bilayers composed of 70% zwitterionic C20 diphytanyl-glycero-phosphatidylcholine lipid and 30% C20 diphytanyl-diglyceride but melimine can interact with these lipid layers [58].

Both melimine and Mel4 could interact with LTA. This interaction is likely due to their positive charge and the negative charge of LTA given by its phosphate groups. Melimine has an overall positive charge of +16, whilst Mel4 has +14 [31], and this small difference may partly explain some of the different effects of their interactions with LTA. At their MICs, there were 3 times more molecules of Mel4 present than melimine yet the inhibition of bacterial growth was not different, perhaps indicating a reduced ability of Mel4 to interact with LTA. In the ELISA assay at their MICs or MBCs melimine neutralized approximately the same number of LTA molecules as Mel4, further indicating a reduced Mel4-LTA interaction. However, this was not the case with the amount of autolysis observed after incubation of *S. aureus* cell supernatants with *Micrococcus* cells. LTA and teichoic acids bind to staphylococci cell wall and regulate the autolysins activity [59, 60], and addition of AMPs liberate autolysins [61].

A cytoplasmic membrane potential is essential for bacterial replication and ATP synthesis [62]. Dissipation of the membrane potential can increase membrane permeability resulting in loss of ATP [63]. Depletion of intracellular ATP can result in loss of viability of bacteria [62]. ATP leakage was associated with increase of 1.5 log_10_ *S. aureus* bacterial death within 2 min with melimine. However, release of ATP did not correlate with bacterial death with Mel4. Melimine reduced bacterial viability in the ATP assay but not in the membrane depolarization assay at same time point. This effect may be due to the presence of 100 mM KCl in the bacterial suspension media of the membrane depolarisation assay. KCl can increase the MIC of AMPs against *S. aureus* [64]. The amount of ATP released by Mel4 plateaued after four minutes exposure and never reached to the level of melimine until 10 minutes exposure. ATP may be depleted in the supernatant of Mel4 treated cells possibly by hydrolysis at their cell surfaces. Mel4 released small amounts of ATP without permeabilizing the cell membrane. ATP being a smaller molecule needs only a pore size of 1.5 nm to leak out from compromised membranes [65]. Thus, pores size produced by Mel4 in the cell membrane are enough for ATP release as they do not need well defined pores in the membrane to escape from bacteria. A comprehensive timeline of the mechanism of action of melimine and Mel4 against *S. aureus* is summarized in Fig. 10.

### Conclusions

It is likely that the amphipathic characteristics of melimine allowed disruption of the cell membranes and pore formation that resulted in ATP and DNA release and ultimately cell death. However, Mel4 showed less interaction with cell membranes and its killing of *S. aureus* was more likely due to activation autolysins along with minimum membrane disruption.

## Acknowledgements

The first author is grateful to Higher Education Commission (HEC) of Pakistan and the University of New South Wales, Australia for provision of PhD scholarship. The authors are also grateful to Christopher Brownlee of the Biological Resources Imaging Lab (BRIL) at the University of New South Wales, Australia for helping in Flow Cytometry analysis.

## Authors Contributions

Conceived and designed the experiments, M.Y., and M.W.; performed the experiments and analysed the data, M.Y.; contributed to the writing of the manuscript and analysis of data; M.Y., and M.W.; edited the manuscript; D. D. All the co-authors read and approved the final manuscript.

## Figure Legends

**Fig. 1** Growth of *S. aureus* in the presence and absence of LTA.. The antibacterial effect of melimine and Mel4 was significantly reduced by addition of LTA (*p*=0.004) after 6 h of incubation.

**Fig. 2** Cytoplasmic membrane depolarization of *S. aureus* by melimine and Mel4, as assessed by release of the membrane potential-sensitive dye DiSC3-(5) measured spectroscopically at 622_nm_ to 670_nm_ excitation and emission wavelengths, and corresponding bacterial survival as determined by plate counts. Data presented as means (±SD) of three independent repeats in triplicate. DMSO used as a positive control and buffer only as a negative control.

**Fig. 3** Cytoplasmic membrane permeability of *S. aureus* by melimine and Mel4 at their MIC and MBC. Fluorescence due to binding of Sytox green with DNA was measured spectroscopically at 480_nm_ to 522_nm_ excitation and emission wavelength. Data presented as means (±SD) of three independent repeats in triplicate. Triton-X 100 used as a positive control and buffer only as a negative control.

**Fig. 4** Membrane permeabilization of *S. aureus* 38 produced by melimine and Mel4 at their MICs as well as Triton X 100 as a positive control and buffer only as a negative control determined by flow cytometry with Syto-9 (membrane permeable) and Propidium Iodide (membrane impermeable stains).

**Fig. 5** The effect of melimine and Mel4 at their MIC and MBC on ATP release from *S. aureus* and the corresponding change in the number of viable cells. Data presented as means (±SD) of three independent repeats in triplicate compared with buffer-treated control.

**Fig. 6** Release of DNA/RNA from *S. aureus* due to action of melimine and Mel4 at their MIC and MBCs determined spectroscopically at OD_260nm._ Data presented as means (±SD) of three independent repeats in triplicate compared with buffer-treated control.

**Fig. 7** Lysis of *S. aureus* by melimine and Mel4 at their MIC and MBC measured spectroscopically at OD_620nm._ Data are shown as means (±SD) of three independent repeats in triplicate compared with buffer-treated control.

**Fig. 8** Killing kinetics of *S. aureus* by melimine and Mel4 at their MIC and MBCs determined spectroscopically at OD_260nm._ Data are presented as means (±SD) of three independent repeats in triplicate compared with buffer-treated control.

**Fig. 9** Release of autolytic activity from *S. aureus* 38 by melimine and Mel4. **(A)** Release of autolysins was monitored by determining the zone of inhibition (arrow) of cell-free supernatants against *Micrococcus lysodeikticus* ATCC 4698 after (I) melimine and (II) Mel4 treatment. 1A = supernatant from melimine treated *S. aureus* 38, IIA supernatant from Mel4 treated *S. aureus*, IB and IIB zones produced by lysozyme (5 mg/ml), IC no zone produced when melimine was incubated with buffer alone, IIC Mel4 incubated with buffer alone, ID and IID *S. aureus* incubated without addition of melimine or Mel4 **(B)** Autolytic effect was also monitored by measuring decrease in OD_570nm_ of PGN suspensions following treatment with cell-free supernatants of melimine and Mel4 treated *S. aureus*. Cell-free supernatants from Mel4 treated *S. aureus* 38 resulted in a greater degradation (≤0.004) of PGN than those from melimine or buffer treated *S. aureus*. Data presented as means (±SD) of three independent repeats in triplicate.

**Fig. 10** Timeline of melimine or Mel4 interacting with *S. aureus*. Both AMPs induced cell membrane depolarization after 30 seconds exposure to the staphylococci followed by release of ATP after 2 minutes. Cell membrane permeabilization occurred with melimine only after 10 minutes with near simultaneous release of DNA/RNA. Mel4 caused transient cell membrane permeability after 30 minutes with no release of DNA/RNA. Complete bacterial lysis started after 6.5 hours of incubation with both the peptides.

